# Focal optogenetic suppression in macaque area MT biases direction discrimination and choice confidence, but only transiently

**DOI:** 10.1101/277251

**Authors:** Christopher R. Fetsch, Naomi N. Odean, Danique Jeurissen, Yasmine El-Shamayleh, Gregory D. Horwitz, Michael N. Shadlen

## Abstract

Insights from causal manipulations of brain activity depend on targeting the spatial and temporal scale most relevant for behavior. Using a sensitive perceptual decision task in monkeys, we examined the effects of randomly-interleaved, rapid, reversible inactivation on a spatial scale previously achieved only with electrical microstimulation. Inactivating neurons in area MT with consistent direction tuning produced systematic effects on choice and confidence. Behavioral effects were attenuated over the course of each session, suggesting compensatory adjustments in the downstream readout of MT over tens of minutes. Compensation also occurred on a sub-second time scale: behavior was largely unaffected on trials with visual stimuli (and concurrent suppression) longer than ∼350ms. These trends were similar for choice and confidence, consistent with the idea of a common mechanism underlying both measures. The findings demonstrate the utility of hyperpolarizing opsins for linking neural population activity at fine spatial and temporal scales to cognitive functions in primates.

## Introduction

To understand how neural activity gives rise to behavior, a powerful approach is to manipulate groups of neurons defined by particular functional or anatomical properties in the context of a suitable behavioral task. This paradigm has grown in popularity over recent years with the advent of sophisticated tools for manipulating neural circuit function. However, recent perspectives have cautioned that so-called causal evidence is not always as decisive as it may appear (Jazayeri and Afraz, 2017), and by itself does not generate the level of understanding we wish to attain (Krakauer et al., 2017). These and other arguments serve to renew a longstanding dictum in systems neuroscience, namely the paramount importance of developing a rigorous theoretical or conceptual framework for understanding the behavior of interest. Only then can clear hypotheses be articulated about the causal role of neural populations or circuits in generating the behavior.

One line of research within the study of perceptual decision making has achieved a degree of progress toward this goal, in part by leveraging detailed knowledge of the neural representation of sensory evidence supporting the decision (reviewed by Newsome, 1997; Cohen and Newsome, 2004; Shadlen and Kiani, 2013). The behavioral task requires judgment of the net direction of motion in a noisy visual display designed to promote the integration of motion information across the display and over time. Neurons in motion-sensitive cortical areas, especially the middle temporal (MT) and medial superior temporal (MST) areas, are well suited to provide the evidence for the task, and theoretical considerations (Shadlen et al., 1996; Gold and Shadlen, 2001) point toward a simple and plausible computation: the subtraction of spike counts (or rates) between pools of neurons favoring each of the direction alternatives. This difference furnishes a quantity proportional to the log likelihood ratio favoring a given alternative, a statistical measure of the weight of evidence that can be accumulated over time to support optimal statistical decision making (Wald, 1947; Gold and Shadlen, 2002).

Building on the work of Newsome and colleagues (Celebrini and Newsome, 1995; Salzman et al., 1992), Ditterich et al. (2003) took advantage of the columnar organization of MT/MST to demonstrate, beyond simply a causal role for these neurons in the task, the specific differencing-and-integration mechanism that was previously hypothesized on theoretical grounds. Using electrical microstimulation (μStim) to activate neurons largely within a single direction column, they confirmed earlier reports showing that monkeys’ choices were biased toward the preferred direction of the activated neurons. More importantly, μStim increased the speed of decisions in favor of the preferred direction while slowing decisions made in favor of the opposite direction. The results were quantitatively consistent with a model in which the decision is formed by temporal integration of momentary evidence defined as the difference in activity between preferred and anti-preferred pools of neurons in MT (Ditterich et al., 2003). More recently, we showed that decision confidence is affected by μStim in a way that is commensurate with the effect on choices and well explained by a bounded evidence accumulation model (Fetsch et al., 2014). Taken together, these findings suggest that a common process of evidence accumulation underlies all three behavioral measures of decision making in this task: choice, reaction time, and confidence.

In general, causal activation (e.g., with μStim) is able to test sufficiency, but not necessity, in the causal chain from neurons to behavior. Thus, reversible inactivation offers an important complement to stimulation, but conventional methods in primate neuroscience (e.g., pharmacological, thermal, or lesion approaches) are lacking in both spatial and temporal specificity. The temporal component is crucial and often neglected; indeed, the insights gained from the μStim studies described above depended on targeting the perturbation not only to the appropriate neurons but also the appropriate time frame (Seidemann et al., 1998). μStim itself has other limitations, including the possibility of antidromic activation, the generation of unwanted temporal and spatial patterns of inactivation in addition to local activation (Butovas and Schwarz, 2003; Logothetis et al., 2010; Seidemann et al., 2002), and the difficulty of quantifying concurrent changes in neural activity due to voltage artifacts.

To address these limitations, we used an optogenetic approach to suppress MT activity during a motion discrimination task with post-decision wagering (PDW; Fig. 1A). While some primate optogenetics studies set out to illuminate the largest possible volume of tissue (Acker et al., 2016; Gerits et al., 2012), we used very low light power levels to target relatively small clusters of excitatory neurons with consistent selectivity for motion. The reasoning is that, in MT, such clusters constitute a critical functional unit for the conversion of a sensory representation into evidence for the decision. We found that optogenetic suppression was capable of inducing a choice bias against the neurons’ preferred direction, and a corresponding change in in the pattern of post-decision wagering, consistent with a common mechanism underlying choice and confidence. We also found intriguing evidence for compensatory changes in the readout of MT activity, on both the sub-second and tens-of-minutes time scales. The results highlight the importance of spatial and temporal specificity in causal interventions, and they suggest a possible explanation for weak behavioral effects of optogenetic manipulations in some previous nonhuman primate studies.

**Figure 1.**
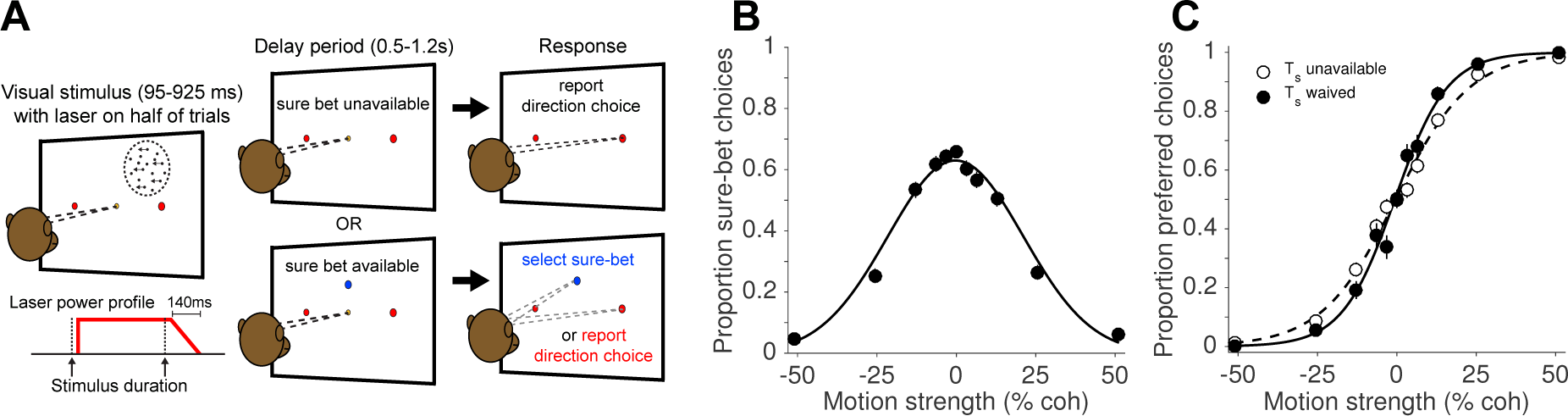
**(A)** Direction discrimination task with post-decision wagering (PDW). The monkey fixates on a central fixation point to initiate the trial. Two red choice targets are presented followed by a random-dot motion (RDM) stimulus in the receptive field of the recorded neurons (left panel). On a random half of trials, a ‘sure-bet’ target (T_s_) is presented (blue spot in the lower panels). After a delay period, the monkey can report his choice by a left-or rightward saccade to one of the red targets to obtain a large juice reward if he is correct, or, if available, choose the sure-bet target for a small but guaranteed reward. On half of trials (randomized independently from T_s_), the RDM stimulus was accompanied by red laser illumination (step-rampdown power profile) of a cluster of neurons expressing the light-sensitive chloride pump Jaws. **(B)** Proportion sure-bet choices in no-laser trials as a function of motion strength (percentage coherently moving dots). **(C)** Proportion of preferred direction choices on no-laser trials as a function of motion strength when the sure bet option was unavailable (dashed) or available but waived (solid).

## Results

We examined the effects of optogenetic inactivation (hereafter, photosuppression) in extrastriate visual cortex (area MT) on perceptual choices and decision confidence. Two monkeys were trained to perform a random-dot motion (RDM) direction discrimination task with post-decision wagering (PDW; Fig. 1A). The task was to decide whether the net direction of dot motion was to the left or to the right, and to indicate the choice with a saccade to a leftward or rightward target when prompted. In addition to the choice targets, a ‘sure bet’ target was presented during the delay period on a random half of trials, allowing the monkey to receive a guaranteed but smaller reward and thereby indicate its lack of confidence in the binary left-right decision. As in previous studies (Fetsch et al., 2014; Kiani and Shadlen, 2009; Zylberberg et al., 2016), monkeys chose the sure bet most frequently when the motion was weak (Fig. 1B) and of short duration (Supplementary Fig. S1B), and their accuracy was greater when the sure bet was available but waived, versus when it was unavailable (Fig. 1C; p < 10^−10^, logistic regression). These behavioral observations, and quantitative analyses published previously (Fetsch et al., 2014; Kiani and Shadlen, 2009), serve to validate the PDW assay as a measure of confidence—that is, a prediction of accuracy based on the state of a decision variable on a given trial, rather than a low-level estimate of trial difficulty or an index of lapses of attention.

### Histological and physiological characterization of Jaws expression

At least 8 weeks before commencing experiments, area MT in one hemisphere of each animal was injected with an AAV vector to drive expression of the red light-sensitive chloride pump Jaws (cruxhalorhodopsin; Chuong et al., 2014) under the control of the CaMKII*α* promoter. Immunohistochemical analysis in a third animal—following injections in a different cortical region (lateral intraparietal area, LIP)—revealed dense Jaws-GFP expression in superficial and deep layers (Fig. 2A), and a tendency to target excitatory pyramidal cells (Fig. 2B), as shown in previous studies using the CaMKII promoter (Han et al., 2009; Nassi et al., 2015). This tendency was not as strong as in previous work: 9% of Jaws-GFP-positive neurons (83 of 903) were double-labeled for the inhibitory marker parvalbumin (Fig. 2B), giving an upper bound on selectivity of 91%, compared to >98% in macaque primary visual cortex (2 of 119 cells double-labeled for any of three different inhibitory markers; Nassi et al., 2015), and 100% in a study of the frontal eye field (0 of 78 cells double-labeled for GABA; Han et al., 2009). However, in addition to being conducted in different cortical areas, these studies used lentiviral vectors rather than AAV. Thus, the effectiveness of promoter-based targeting of excitatory neurons likely depends on the viral vector and/or serotype, and could also vary across brain areas.

**Figure 2.**
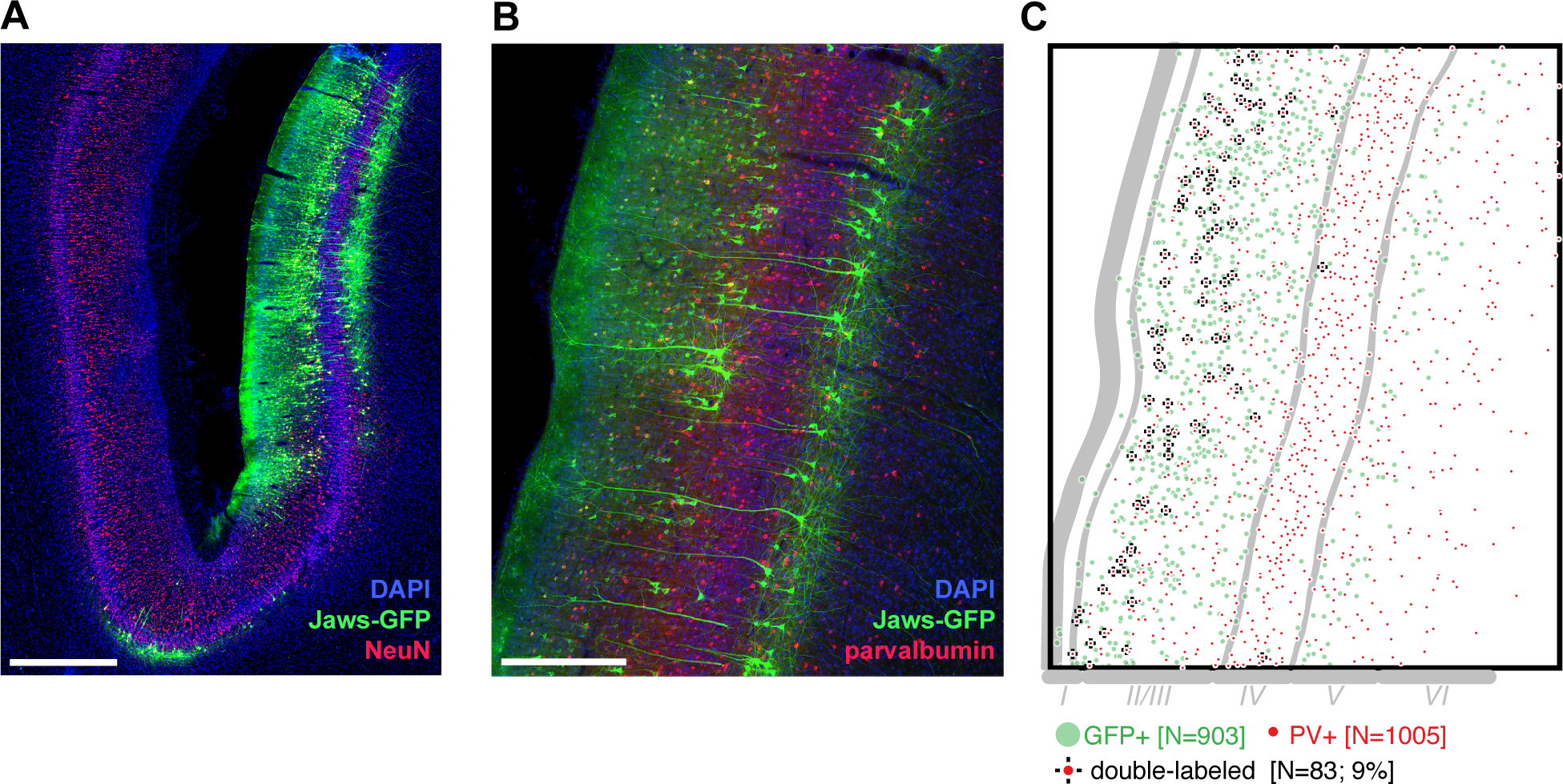
**(A)** Histological section imaged at 10X showing expression of Jaws (green) following a series of viral injections along a single injectrode penetration in lateral intraparietal cortex (LIP). Laminar distribution, aided by visualizing NeuN immunoreactivity (red), appears concentrated in layers II/III and V. Blue: DAPI. Scale bar = 1 mm. **(B)** 20X image of a different section stained for Jaws-GFP (green) and the inhibitory marker parvalbumin (PV, red). **(C)** Cell counts from the image in B, quantifying the small but not insignificant double-labeling of Jaws-GFP and PV. Scale bar = 500 μm.

In each experimental session, we advanced a custom optrode into area MT and began searching for suitable sites for testing the effects of targeted photosuppression on behavior. If a suitable site was found (see Materials and methods), the monkey commenced the task described above, with low-power red illumination (633 nm, total power = 0.25-2.0 mW, irradiance = 2-16 mW/mm^2^ at a distance of 300 μm) delivered concurrently with visual stimulation on a random half of trials. Note that unlike previous studies, a red-shifted opsin was chosen not for its suitability for large-volume tissue illumination (Acker et al., 2016) but for its superior photocurrent at very low irradiance (Chuong et al., 2014). Optrode design, laser power, and site selection criteria were all intended to limit the light-induced suppression of activity to a relatively small cluster of MT neurons with consistent receptive field and tuning properties (i.e., preferred speed and direction of motion).

While mapping potential sites for photosuppression based on multi-unit (MU) activity, we occasionally isolated single neurons and used this opportunity to better quantify the physiological effects of the light. Single-unit visual responses to high-coherence RDM stimuli—presented at the preferred direction and speed—were strongly suppressed on average (Fig. 3A), although a handful of neurons showed no effect or an increase in firing rate. Presumably this includes cells that did not express Jaws, as well as a small minority that were indirectly activated through polysynaptic mechanisms (i.e., disinhibition). Among putative Jaws-expressing neurons, the visually evoked response, after subtracting baseline activity, was suppressed by 101% (±13% SEM; N=20; Fig. 3B). Without baseline subtraction, the average firing rate during stimulus presentation was reduced by 72% ± 5%. In contrast, MU responses during the discrimination task were reduced by an average of 33% (± 0.3% SEM, N=20058 trials; Fig. 3C). The fractional degree of suppression (Equation 4; Materials and methods), calculated from MU activity, varied substantially both within and across sessions (Fig. 3A, inset; Fig. 4A). We take this this variability into consideration in the analyses to follow.

**Figure 3.**
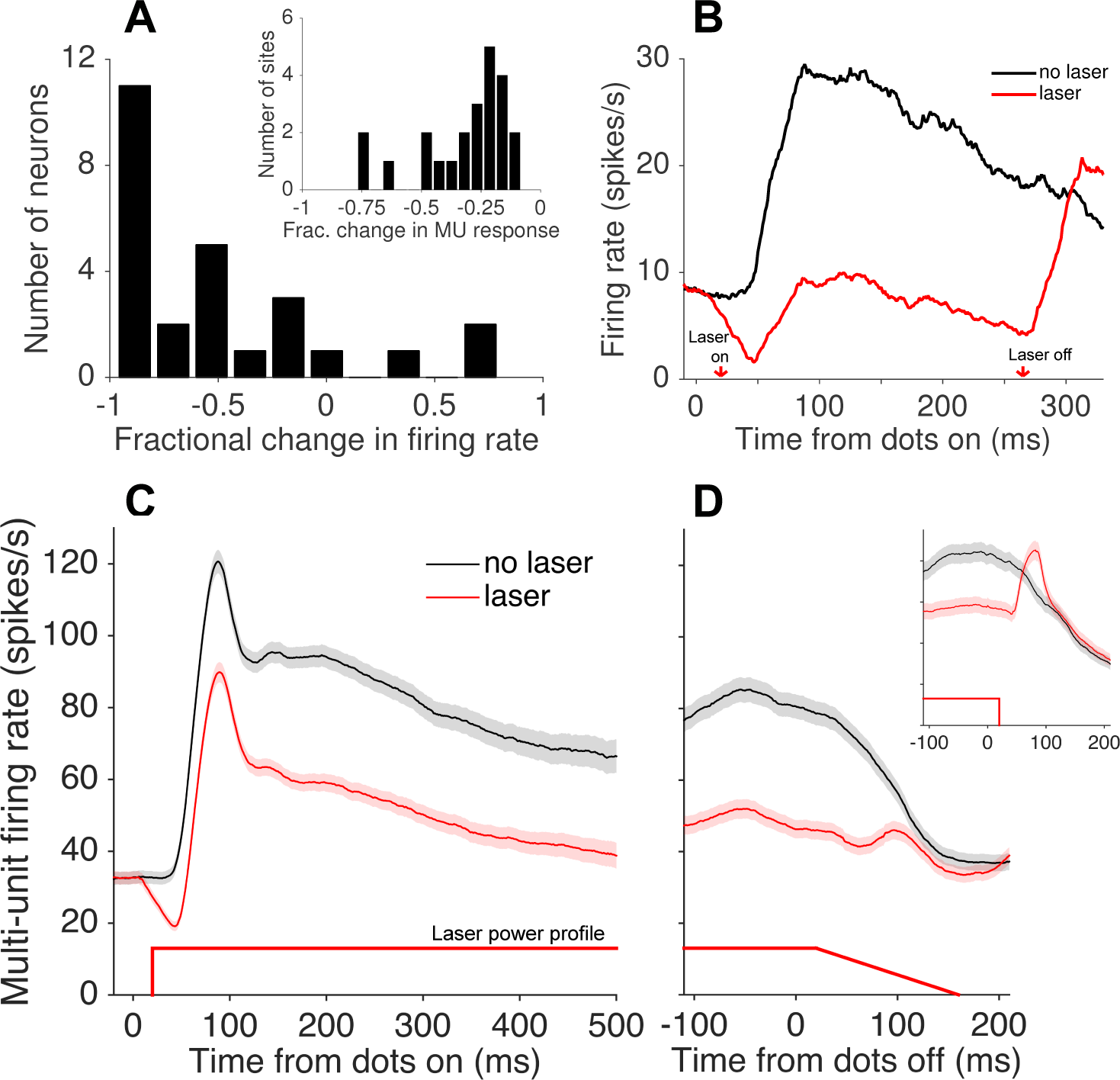
**(A)** Fractional change in firing rate (Δ*R*) of isolated single-units (N=26) in response to a high-coherence RDM stimulus and laser suppression, relative to RDM stimulus alone. The inset shows the fractional change in multi-unit activity recorded at the 23 sites tested behaviorally. **(B)** Average firing rate of single units showing significant Jaws-mediated suppression (N=20). Laser onset occurred 20 ms after stimulus onset, resulting in suppression that preceded the onset of visually-driven activity (i.e., firing rate driven below baseline before recovering to near-baseline levels during visual stimulation). A post-suppression rebound of activity was observed after turning the laser (and visual stimulus) off. Traces depict spike counts in 1-ms bins convolved with a 40-ms causal boxcar filter **(C, D)** Average multi-unit activity for N=23 sites passing the selection criteria for the behavioral task (see Materials and methods), aligned to stimulus (dots) onset **(C)** and offset **(D)**. The majority of sessions included a ramp-down of 140 ms in laser power starting 20ms after stimulus offset, reducing the post-suppression burst seen when no down-ramp was used (inset). Shaded region shows ± SEM.

**Figure 4.**
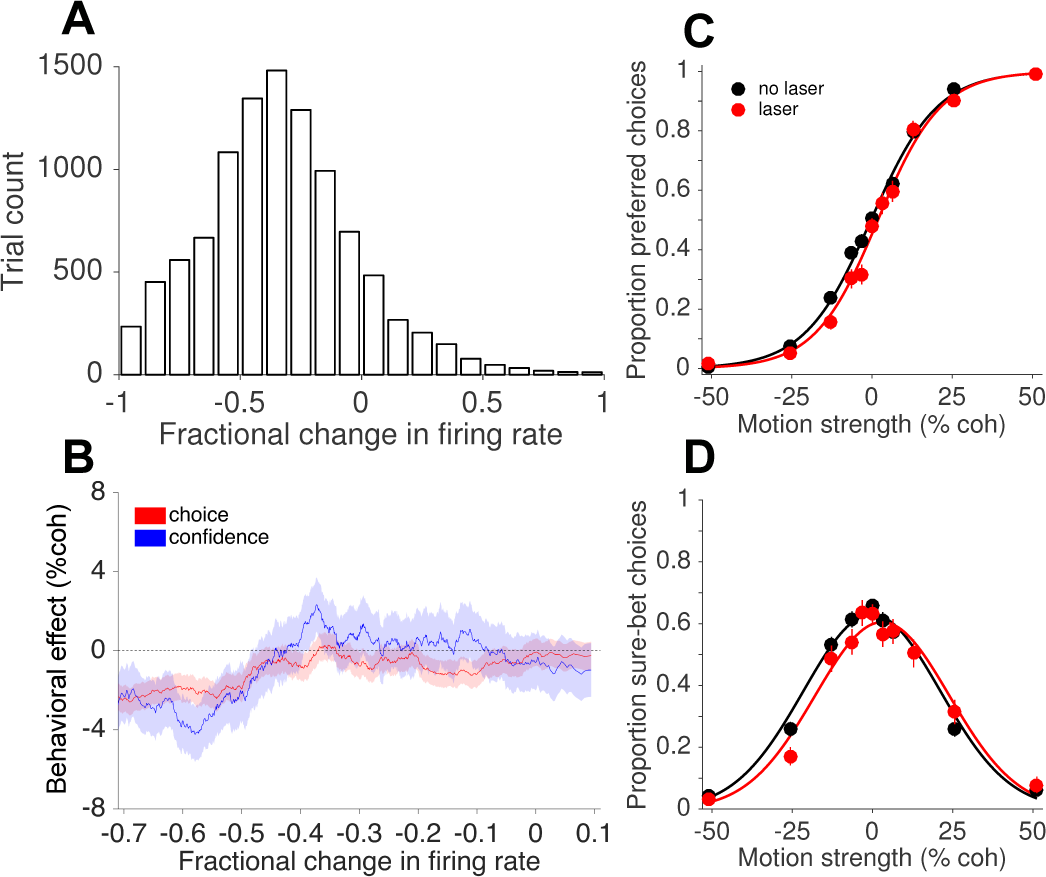
**(A)** Histogram of the fractional change in multi-unit firing rate (Δ*R*), relative to the mean of no-laser trials for the corresponding session and trial type (Equation 4). Across-trial variability is high, allowing for an analysis of how the degree of suppression relates to changes in behavior. **(B)** Behavioral effects, expressed as the horizontal shift in the logistic choice function (Equation 1, red) and bell-shaped confidence function (Equation 3, blue), as a function of fractional change in firing rate. Shaded error regions indicate ± SEM. **(C)** Choice functions for the set of laser trials (red) exhibiting strong suppression (Δ*R* < −0.45, N=3636 trials) compared to all no-laser trials (black, N = 9912 trials). **(D)** The corresponding confidence functions (proportion sure-bet choices as a function of motion strength) for the set of trials shown in C.

Note that the expected rebound of spiking after abrupt laser offset (Fig. 3B) was mitigated by introducing a 140-ms ramp-down of laser power in the majority of behavioral sessions (Fig. 3D; Chuong et al., 2014). We did not detect a difference in behavioral effects when the rebound was suppressed, compared to an earlier subset of sessions without the ramp-down (Fig. 3D, inset), and therefore pooled all sessions for subsequent analyses.

### Behavioral effects of photosuppression

To introduce the behavioral analyses presented below, we first revisit results from recent μStim experiments using the same behavioral task (Fetsch et al., 2014). Supplementary Figure S2 shows μStim data from one of the two monkeys used in the previous study who also participated in the present study. Microstimulation in areas MT and MST altered choice and confidence in a manner that largely mimics a change in the motion strength, with a directionality predicted by the preferred direction of the stimulated neurons. Although there were modest changes in the slope of the choice function and height of the confidence function, here for simplicity we focus on the lateral shift of the functions in units of motion strength (% coherence; Equations 1 and 3, Materials and methods). The direction and size of these shifts was well matched in the aggregate, and highly correlated across sessions, suggesting that MT/MST activity contributes to a common decision variable that governs both choice and confidence (Fetsch et al., 2014).

Under this interpretation, suppression of MT activity—provided it is targeted to a population of neurons with consistent tuning—should cause shifts in the opposite direction: fewer choices in the preferred direction, and a rightward shift of the bell-shaped confidence function. Moreover, one would expect behavioral effects to depend on the degree of suppression, which as noted above, was highly variable. For all laser trials in each session, we calculated the fractional change in multi-unit firing rate (Δ*R*; Equation 4) relative to the mean of no-laser trials for the corresponding session and trial type (motion direction and coherence; Fig. 4A). We then estimated the behavioral effect size (Equations 1 and 3) as a function of neuronal effect size, using a sliding window of trials pooled across sessions and sorted by Δ*R*. This analysis (Fig. 4B) showed that photosuppression caused a modest but significant bias in choice and confidence in the predicted direction, provided that the degree of response suppression was greater than about 45% (choice shift = −2.3% coh ± 0.6% SE, N = 13548 trials with Δ*R* < −0.45; p = 0.0001; Fig. 4C; confidence shift = −3.6% ± 1.2%, p = 0.004; Fig. 4D). Notably, the effects (or lack thereof) on choice and confidence were reasonably well matched across the full range of Δ*R*, supporting the idea of a common decision mechanism linking the two measures.

To account for potential contributions of neuronal response variability independent of the variation in degree of suppression, we performed the following control analysis. Taking only no-laser trials, we randomly assigned 50% of them a fictitious category (“sham laser-on trials”), sorted these trials as above by their fractional difference relative to the mean for a given session and trial type (Supplementary Fig. S3A), and performed the same sliding-window analysis of choice bias and shifts of the confidence functions. We then repeated the procedure 100 times with different randomized trial assignments and averaged the results. The resulting traces (Supplementary Fig. S3B) showed a weak trend consistent with a covariation between response fluctuations and behavior (i.e., choice probability; Britten et al., 1996), but which was severalfold smaller than in the original analysis that included actual photosuppression (Fig. 4B) and not statistically significant (p = 0.42, Equation 5, *β*_6_). This implies that the behavioral effects we observed under strong photosuppression are not explained away by normal response variability and its covariation with choice (and confidence).

### Temporal factors influence behavioral effects: compensation on multiple time scales?

The result depicted in Figure 4B implies that photosuppression was only effective at altering behavior for a subset of trials. Indeed, most individual sessions failed to show statistically reliable effects on behavior when analyzed in their entirety (Supplementary Fig. S4). However, while collecting the data we noticed a tendency for systematic biases on laser trials to appear early within a session, only to dissipate over the course of tens of minutes (100-500 trials; ∼5 s per trial on average, including intertrial intervals). This can be seen in the example session shown in Figure 5. Over the first 500 trials, the monkey tended to choose the preferred direction less often on laser trials vs. no-laser trials, effectively shifting the psychometric curve to the right (Fig. 5A; shift = −4.6%, p = 0.14, Equation 1). In contrast, there was essentially no effect on choices in the remaining ∼1000 trials of the session (Fig. 5B; shift = −0.7% coh, p = 0.75). The pattern was similar for the confidence assay: a rightward shift of the confidence function on early trials (Fig. 5C; shift = −4.2%, p = 0.34) but not later trials (Fig. 5D, 1.2%, p = 0.80). This trend was evident when pooling across all sessions with enough trials to measure it (minimum 800 trials per session, N=10 sessions): rightward shifts early in the session (choice shift = −3.5% coh ± 1.1%, p = 0.001, Fig. 5A’; confidence shift = −3.8% ± 1.9% [Equation 3], p = 0.05, Fig. 5C’) but not later (choice shift = −0.7%, p = 0.26, Fig. 5B’; confidence shift = 0.1%, p = 0.93; Fig. 5D’). Post-hoc tests using generalized linear models (Equation 5, see Materials and methods) showed that the difference between early and late trials was statistically reliable for choice (p = 0.01) and confidence (p = 0.02, Bonferroni corrected).

**Figure 5.**
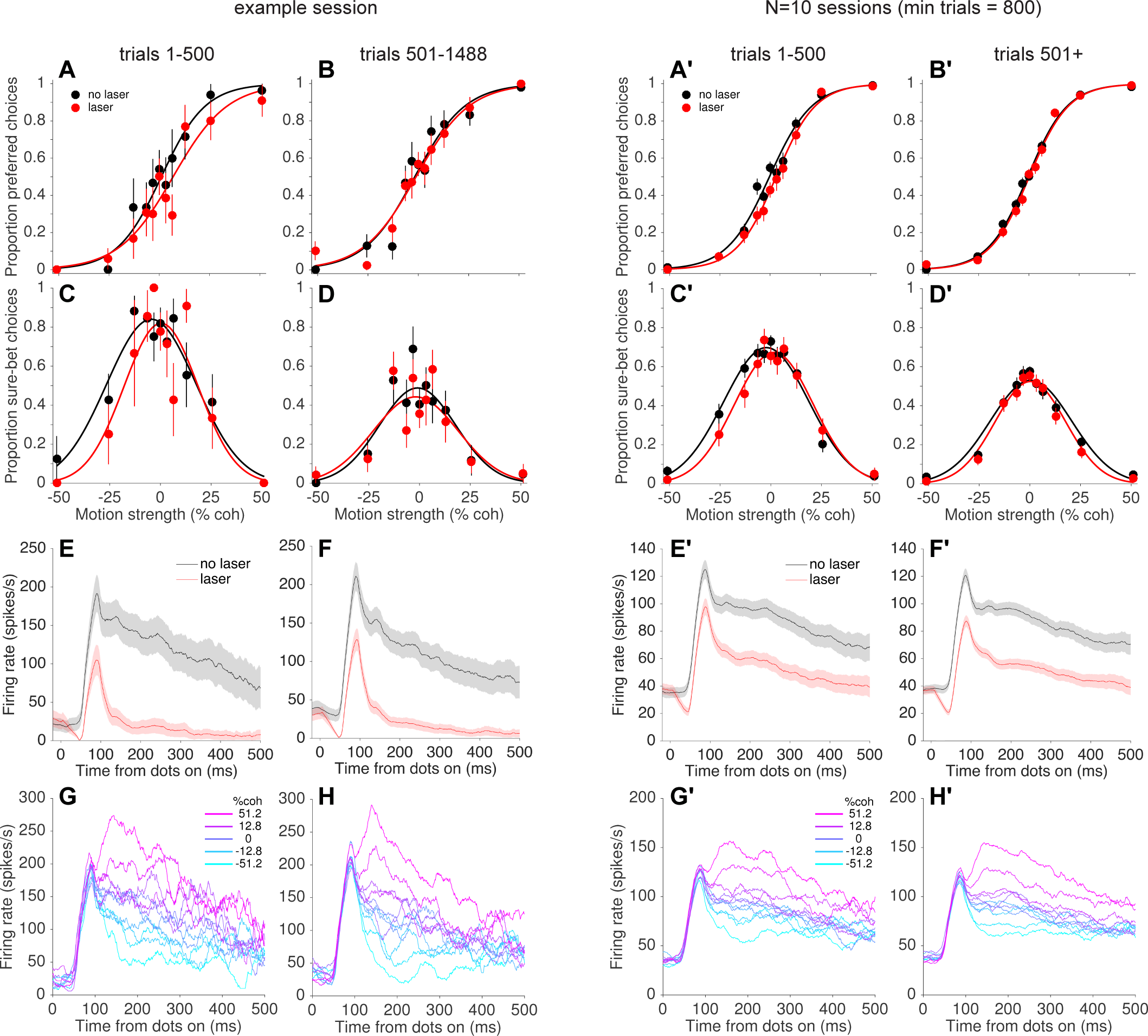
Comparison of early and late trials in an example session (5A-H) and for all sessions with >800 trials (5A’-H’, N=10 sessions, 10927 trials with Δ*R* < −0.25). Systematic behavioral effects of photosuppression can be seen in the early but not the late trials, whereas suppression of activity and direction selectivity remain largely unchanged. **(A,B)** Choice functions for laser (red) and no-laser (black) trials early in the session (A, trials 1-500) versus late (B, trial number > 500). **(C,D)** corresponding confidence functions for the same trials in A and B, respectively. The decrease in overall sure-bet proportion between C and D, indicating an increase in overall confidence over the course of a session, was a behavioral peculiarity of one monkey that was unrelated to photosuppression (i.e., occurred throughout training and in no-laser control sessions). **(E,F)** Average multi-unit activity showing degree of suppression for early (E) and late (F) trials. **(G,H)** Average firing rate on no-laser trials, early (G) vs. late (H) in the session, separated by signed motion coherence (positive = preferred direction, magenta; negative = null direction, cyan).

Salzman and colleagues (1992) reported that the effects of μStim dissipated over the course of the behavioral sessions, which they interpreted as indicating neuronal fatigue, damage, or shifts in the position of the stimulating electrode. In our case, however, the decrease in effectiveness of photosuppression on behavior was not accompanied by a pronounced decrease in its effect on neural responses (compare separation between black and red curves in Fig. 5E versus Fig. 5F, and 5E’ versus 5F’ for the pooled dataset), nor by a qualitative change in the direction selectivity of the affected neurons (Fig. 5G vs. 5H, and 5G’ vs. 5H’). At the conclusion of this and several other sessions, we re-measured direction tuning at multiple depths and found little evidence for optrode drift, tissue damage or response fatigue. Instead, the results suggest a compensatory change in the downstream readout of MT neural signals to reduce or nullify the effects of photosuppression on a time scale of roughly 10-40 minutes.

To examine this putative compensation at a finer grain, we plotted the choice and confidence effects calculated in a sliding window of trials sorted by trial number after pooling across sessions (Fig. 6A; N = 8057 trials from 23 sessions; see Materials and methods). Interestingly, the effect on confidence (blue trace) appears to dissipate more quickly than the effect on choice, and even @’overshoots’ zero for a period of time roughly between trial numbers 500-900 (∼40-75 min), before settling back to zero later in the session. Recall that positive values on the ordinate indicate leftward shifts of the function, as observed previously with μStim, and are thus opposite of the expected effect from inactivation. We speculate on the reasons for this puzzling pattern in the Discussion. Lastly, for comparison with previous work, we repeated this analysis with the μStim dataset depicted in Supplementary Fig. S2 (Fetsch et al., 2014). Unlike photosuppression, and contrary to the findings of Salzman et al. (1992), the biases in choice and confidence induced by μStim remained relatively stable throughout the behavioral session (Supplementary Fig. S5A). This result implies that the capacity of cortical circuits to compensate for artificially induced activity patterns may be greater for photosuppression compared to μStim, although we cannot ascertain whether the key difference is the sign (suppression versus stimulation) or the modality (optogenetic versus electrical) of the perturbation.

**Figure 6.**
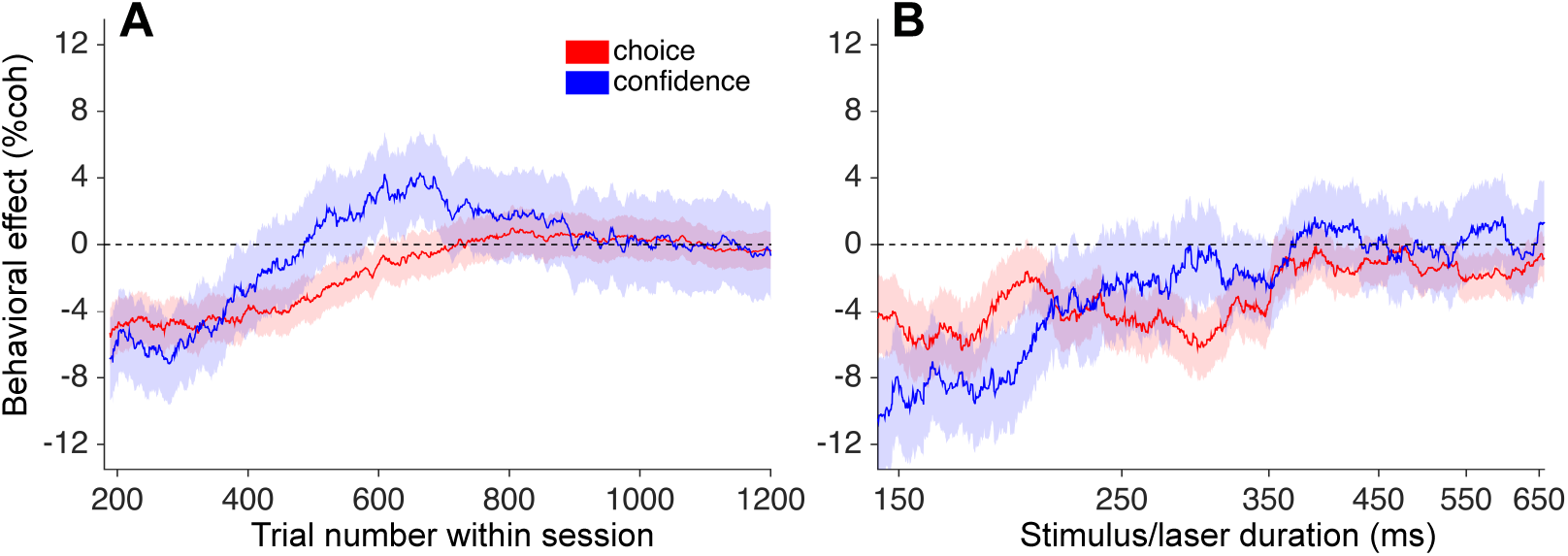
Attenuation of behavioral effects on long and short time scales. **(A)** Shifts of the choice (red) and confidence (blue) curves (± SEM) as a function of trial number (∼8 minutes per 100 trials). Trials were pooled across sessions, conditioned on Δ*R* < −0.25 and duration < 300 ms (N=8057), and behavioral effects (Equations 1 and 3) were calculated over a sliding window of ∼1750 trials sorted by trial number within session. **(B)** Corresponding sliding-window analysis of trials sorted by stimulus/laser duration, after conditioning on Δ*R* < −0.25 and trial number < 500. N = 7381 trials, window width = 1600 trials.

The presence of compensatory changes in readout across trials raises the question of whether such changes could occur on a faster time scale as well. We addressed this question by grouping trials according to the experimenter-controlled stimulus (and laser) duration, which varied randomly across trials from 95-925 ms (Supplementary Fig. S1A). Remarkably, we found that the behavioral effects of photosuppression were largely restricted to short-and intermediate-duration trials (duration < ∼350 ms; Fig. 6B). Figure 7 depicts this trend pooled across all sessions, in the same format as Figure 5. Short-duration trials exhibited light-induced shifts of −4.7% (p = 0.0006, Fig. 7A) and −5.7% (p = 0.004, Fig. 7C) for choice and confidence respectively, whereas long-duration trials showed no systematic biases in either measure (choice: −0.6%, p = 0.50; confidence: 0.38%, p = 0.84). The effect of duration was statistically significant (Equation 5, Materials and methods) for confidence (p = 0.002, Bonferroni corrected) and marginally so for choice (p = 0.04)

**Figure 7.**
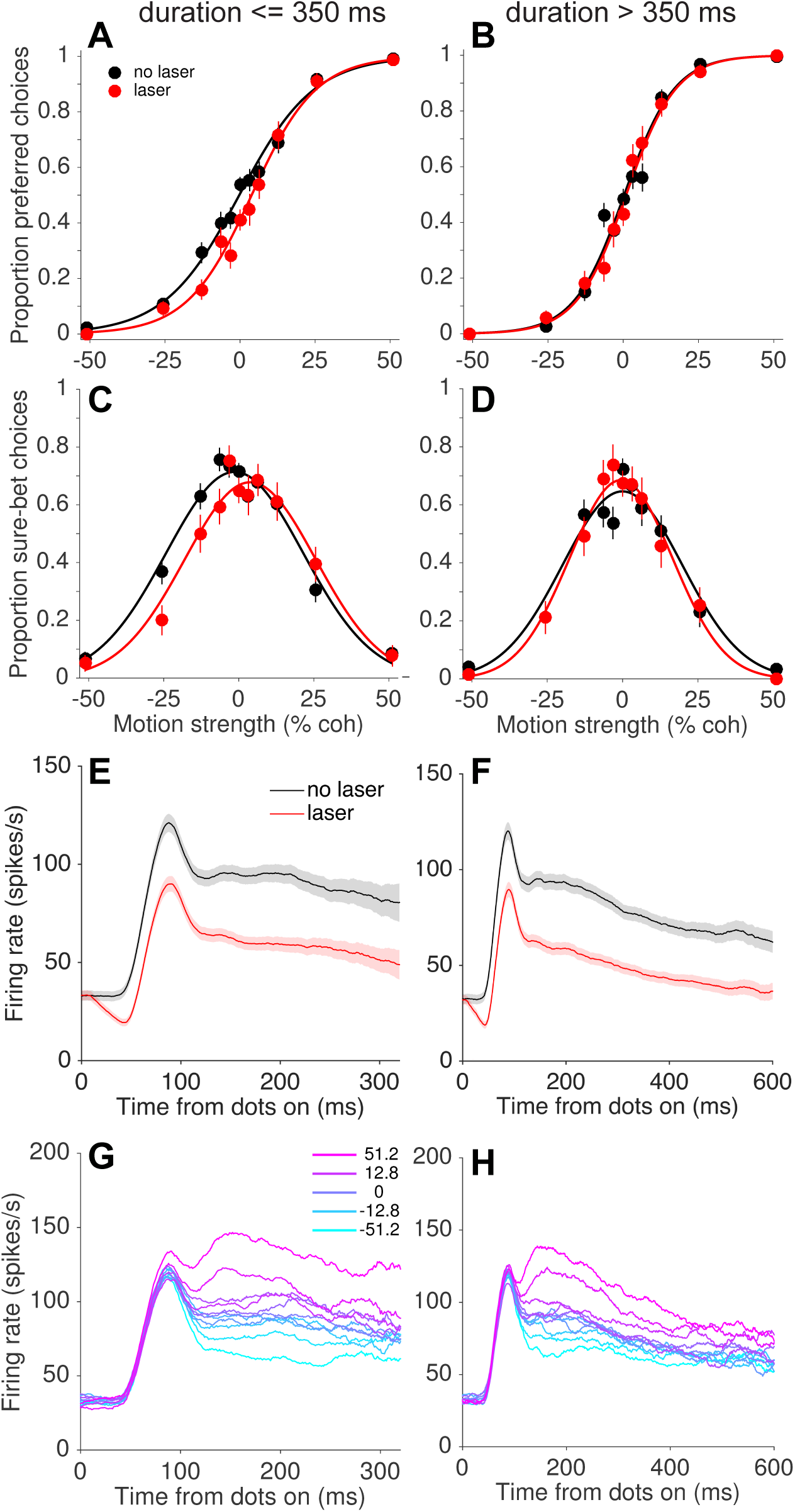
Behavioral effects of photosuppression were present on trials with stimulus/laser epochs of short duration (left column, duration <= 350 ms) but not longer durations (right column, duration > 350 ms), whereas degree of suppression and direction selectivity were similar across durations. Format as in Fig. 5. Data include trials conditioned on Δ*R* < −0.25 and trial number < 500 (N=4249 and 3132 trials for left and right columns, respectively).

As with the slower form of compensation, this sub-second attenuation of behavioral effects was not readily explained by differences in MT neural responses on short-versus long-duration trials (Fig. 7E-H). On long trials, the separation of mean firing rates across motion strengths and directions became smaller over the course of the stimulus epoch (Fig. 7H). One might wonder whether this can explain the behavioral difference, under the assumption that neural signals contribute to behavior in proportion to their sensitivity (Gu et al., 2008; Purushothaman and Bradley, 2005). However, a quantitative analysis incorporating response variability revealed that direction selectivity was actually stronger for long-duration trials, when taking into account spiking activity throughout the trial (lower neuronal threshold on long trials [27.2% coh ± 1.6 SE] compared to short trials [37.5 %coh +/− 2.5 SE]). Thus, the differences in firing rates on the right side of Fig. 7H versus Fig. 7G cannot alone account for the inability of photosuppression to affect behavior on long-duration trials. Indeed, any candidate explanation for this result must contend with the fact that activity early in the stimulus/laser epoch was suppressed equally on short-versus long-duration trials, yet only the former showed behavioral effects. It seems improbable that downstream circuits could treat such early activity differently, because at any given time during a trial there was no way to predict whether the stimulus (and laser) would terminate in the next moment or continue. Below we discuss possible explanations and implications of this surprising finding.

## Discussion

### Focal optogenetic suppression

The results demonstrate that systematic behavioral effects of optogenetic manipulation in nonhuman primates are achievable even when this requires targeting of neural populations on a small spatial scale, that of cortical columns or clusters of neurons with consistent functional properties. This is the scale at which electrical microstimulation has proven most effective, but primate systems neuroscience has lacked an inactivation counterpart with comparable spatial and temporal specificity, relying instead on pharmacological and cooling methods that last minutes to days and affect larger swaths of tissue. There is understandable excitement surrounding the development of optogenetic methods for targeting particular cell types or projections (El-Shamayleh et al., 2017; Klein et al., 2016; Stauffer et al., 2016; Ohayon & DiCarlo, Soc. Neurosci. Abs. 2017), and many questions will require such an approach. Our study shows that another form of targeting—that of spatially segregated sub-populations or circuit components—is possible in monkeys using off-the-shelf viral vectors combined with careful site selection and light delivery.

### A common mechanism for choice and confidence

Decision confidence is defined as the subjective degree of belief that one has chosen the correct or more rewarding option among alternatives. In many natural situations, the decision maker does not receive immediate feedback on the accuracy of a choice, but instead must rely on confidence to guide subsequent decisions that depend on previous outcomes. Confidence also has a powerful influence on learning and adapting to new environments. For instance, when negative feedback follows a choice made with high confidence, it implies that something about the world has changed, triggering a shift in behavioral strategy or increase in learning rate (Dayan et al., 2000; Kepecs, 2013; Purcell and Kiani, 2016).

The neural basis of assigning confidence in a decision is not as well understood as other features of the decision, such as the choice itself and the amount of time it takes to decide. One hypothesis is that the evidence that supports a choice also guides the assessment of reliability of the evidence, hence the probability that the choice is correct. In this vein, Kiani & Shadlen (2009) demonstrated how choice and confidence can arise through a common mechanism of evidence accumulation, and several studies have since provided additional support for this idea (Fetsch et al., 2014; Kiani et al., 2014; van den Berg et al., 2016; Zylberberg et al., 2016). Using the same task as in the present study, we found that the quantitative link between choice and confidence was preserved under MT/MST microstimulation (Fetsch et al., 2014). Changes in both measures—and their specific relationship with motion strength and viewing duration—evinced a commensurate change in the strength of momentary evidence, and the improvement in discrimination sensitivity when confidence was high (Fig. 1C) was retained, as predicted by a bounded evidence accumulation model.

The present data provides an inactivation counterpart to this: photosuppresion in MT biased the animal’s choices away from the preferred direction, and also altered confidence in a way that similarly mimics a change in motion strength (i.e., a rightward shift of the bell-shaped curve describing sure-bet choices as a function of coherence). These effects were well matched as a function of the degree of suppression (Fig. 4B), and they dissipated over a similar time course, both across (Fig. 6A) and within trials (Fig. 6B). These observations are consistent with a role for these neurons in providing a shared source of evidence supporting both aspects of a decision. We did not attempt to fit the bounded accumulation model to the present dataset, in part because of limited sample size after accounting for the factors that influence the behavioral effectiveness of the perturbation (trial number, duration, and variability in the degree of suppression). However, knowledge of these factors, along with ongoing developments in causal tools, could provide an opportunity to deepen our understanding of the neural basis of confidence judgments by exposing deviations from the current model.

In fact, we did observe some exceptions to the coupling between choice and confidence that could be worth further inquiry. The more rapid recovery and overshoot of the confidence effect as a function of trial number (Fig. 6A, blue; see below) was notably absent in the measurement of choice bias (red). The largest effect on confidence, observed at the shortest durations, was greater than the choice bias observed on those same trials (Fig. 6B; but see Fig. S5 to compare to the size of deviations observed with μStim). On a session by session basis, the shifts in the two functions were not significantly correlated (Fig. S4) as they were for μStim (Fetsch et al., 2014; their Fig. S1A), although single-session estimates in the present study were less reliable owing to limited sample size in several sessions. Still, it could be that the processing of sensory evidence that gives rise to choice and confidence utilizes slightly non-overlapping neural circuitry. To explore the anatomical and functional basis of this divergence, future work could combine multi-site recording with manipulations (either stimulus-based or via causal methods) that systematically break the correspondence between choice and confidence. Many recent studies provide examples of related dissociations, and point toward specific frontal lobe and subcortical structures that may be involved (Cortese et al., 2016; Fleming et al., 2012; 2014; Odegaard et al., 2018; Rounis et al., 2010).

### Compensation on slow time scales

Behavioral effects of photosuppression were strongly attenuated over the course of the session (Figs. 5, 6A). This trend was not explained by differences in the physiological effectiveness of photosuppression, nor in the responsiveness or selectivity of the affected neurons. Rather, the results suggest some form of compensation or down-weighting of MT signals by downstream areas that are reading out this activity to form a decision.

Because the biasing effect of photosuppression (as with μStim) increases the error rate on these trials, a compensatory mechanism that counteracts this bias could be advantageous for the animal in terms of reward rate. Evidence for a slow drift in bias or decision criterion comes from the so-called ‘null-choice bias’ observed in μStim experiments (Salzman, et al., 1992; Fetsch et al, 2014). This refers to the tendency of the animal to gradually adopt an overall bias away from the preferred direction of the stimulated neurons, as if to counteract the excess of choices made in the preferred direction as a result of the stimulation. Crucially, however, the null-choice bias is applied to both μStim and no-μStim trials, leaving the overall effect of μStim intact (Supplementary Fig. S2A). In contrast, a compensatory mechanism that permits downstream structures to down-weight the artificially suppressed neural signals could nullify the effects of photosuppresion over time (Fig. 5A’ and 5B’). Since laser trials were randomly interleaved, such a mechanism would require that readout circuitry in some way ‘detects’ the presence of photosuppression on a trial by trial basis, unlike μStim. This does not necessarily imply that the monkey himself detects the perturbation or perceives something anomalous on those trials; rather it could be that downstream circuits automatically learn to take into account signals that are deemed less reliable by virtue of their association with a lower rate of reward.

For confidence (Fig. 6A, blue), the pattern of effects across trials appeared to return to an unbiased state more rapidly than for choice (Fig. 6A, red), and it even showed a transient shift in the opposite direction (leftward, as in Fig. S2B). If the compensation is implemented by a change in the influence of particular MT signals on downstream neurons (the ‘readout weights’), then an overshoot would imply that the weights not only fall to zero but actually reverse sign. The fact that this occurs for confidence but not for choice suggests that the way sensory signals are pooled to construct a degree of confidence may be more flexibly linked to recent outcomes. It could also reflect greater sensitivity—to the point of ‘overreaction’—on the part of the readout mechanism in response to unnatural patterns of activity induced by photosuppression. For now this remains largely speculative, and we should note that caution is warranted given the limited sample size and noisiness of the measurement (difference between means of two Gaussian fits). If this subtle effect is confirmed with additional data, it could be probed further by modifying the behavioral assay to elicit a continuous estimate of confidence on individual trials (Kiani et al., 2014; Lak et al., 2014). Given the broad importance of the balance between excitation and inhibition, it may also be of interest to ask whether the presence and pattern of compensation depends on the cell types being targeted, for instance suppression of excitatory neurons versus activation of inhibitory neurons, or global suppression using a pan-neuronal promoter.

### Compensation on fast time scales

The inability of photosuppression to influence behavior on long-duration trials (Fig. 6B, 7) is surprising, as it implies the presence of a compensatory mechanism operating on a sub-second time scale. Note that this result is not a trivial consequence of the brain placing greater weight on sensory evidence arriving earlier within a trial. Selective influence of early versus late evidence has been observed previously and is often attributed to a bounded decision process (Kiani et al., 2008; Raposo et al., 2012): late evidence is more likely to arrive after the threshold for decision termination has been reached, and therefore has less impact on choices on average. Here, suppression on long-duration trials affects early evidence to the same degree as on short-duration trials (Figure. 7F vs. 7E), yet the former showed virtually no behavioral effects (Fig. 7B+D vs. 7A+C). Selective down-weighting of evidence on long-duration trials is also unlikely to involve slow plasticity mechanisms, because both the presence of illumination and the duration itself were randomized. Instead it seems to require a fast rerouting of information through parallel circuits, effectively enabling the consultation of different populations of neurons from trial to trial. These redundant populations could be within MT, or in other structures such as MST that similarly represent motion evidence for this task.

An intriguing (and not mutually exclusive) possibility is that fast compensation involves neuromodulators such as acetylcholine (ACh). ACh is involved in the regulation of visual processing and attention (Sarter et al., 2005). Within area MT, ACh receptors are expressed in most parvalbumin-positive inhibitory interneurons (Disney et al., 2014), which were largely spared by our CaMKII-based approach (Fig. 2B-C). Although ACh-releasing axons extend across relatively large regions of cortex, it has been proposed that structural features of cortical neuropil—combined with the distribution of ACh receptors and choline transporters—could effectively compartmentalize cortical subregions to permit neuromodulation on much smaller spatial scales than is generally appreciated (Coppola et al., 2016). Moreover, the time scale of muscarinic (G-protein-coupled) ACh receptor activation, albeit slower than ion channel receptors, is potentially within the right range to affect information processing within individual trials (Lohse et al., 2009). Thus, ACh-mediated regulation of local inhibition could conceivably play a role in the fast compensation we observed, although the specifics await additional theoretical and experimental investigation.

## Conclusion

The approach taken in this study was motivated by a large body of research linking the functional properties of sensory neurons to the formation of a perceptual decision. In nonhuman primates, the effectiveness of modern causal tools for manipulating behavior, although not without successes, has been limited relative to other animal models. One sensible response to this reality is to develop methods for illuminating larger volumes of tissue (Gerits et al., 2012; Acker et al., 2016), under the assumption that modulation of primate behavior with optogenetics is hindered by their larger brain size. As an alternative, we suggest that a more nuanced understanding of the relationship between neural populations and behavior can yield insights that would be obscured by the large-volume approach. Naturally, the ideal spatial scale depends not only on the anatomy but on the specific question being addressed. For researchers seeking to exploit the temporal control afforded by optogenetic inactivation, the present findings should motivate a renewed focus on developing sensitive and well-characterized behavioral paradigms, followed by a specific hypothesis-driven account of neural activity before designing an appropriate causal manipulation.

## Materials and methods

### Behavioral task

Two adult male rhesus monkeys (*Macaca mulatta*) performed a direction discrimination task with post-decision wagering (PDW, Figure 1A), as described previously (Kiani and Shadlen, 2009; Fetsch et al., 2014, Zylberberg et al., 2016). In brief, animals were required to fixate a central target, after which two direction-choice targets appeared 9° to the left and right of the fixation point, followed by a dynamic random-dot motion (RDM) stimulus. Properties of the RDM (patch size, position, and dot speed) were set for each experimental session to match the aggregate receptive field (RF) and tuning properties of neurons throughout a 200-400 μm region near the optrode tip (see below for site selection criteria). Owing to behavioral limitations related to the training history of the animals, the net direction of motion was constrained to be predominantly left or right; sites with preferred directions >30° from horizontal were bypassed. Motion strength or coherence (percent coherently moving dots) on each trial was sampled uniformly from the set {0, 3.2, 6.4, 12.8, 25.6, 51.2}, and stimulus duration was drawn from a truncated exponential distribution with range = 95-925 ms, mean = 350 ms, and median = 300 ms.

After motion offset, the monkey maintained fixation through a variable delay period during which a third target (the sure-bet target, T_s_) was presented on a random half of trials. T_s_ differed in color and size from the direction choice targets, and was positioned 6° above the fixation point, perpendicular to the axis of motion and the direction-choice targets. Whether or not T_s_ was presented, the delay period ended with disappearance of the fixation point which acted as a‘go’ cue for the monkey to saccade to one of the targets. If T_s_ was available, the monkey could choose it and receive a guaranteed reward (drop of water or juice), or waive T_s_ and make the higher-stakes direction choice; otherwise only the binary direction choice was available. Correct direction choices yielded a larger liquid reward than T_s_ choices, while errors resulted in a 4.5 s timeout. The ratio of sure-bet reward size to direction-choice reward size (mean = 0.51 for monkey D, 0.62 for monkey N) was set and periodically adjusted to encourage the animals to choose T_s_ approximately 60% of the time at the weakest motion strengths.

### Surgery and injection of viral vector

Procedures were in accordance with the Public Health Service Policy on Humane Care and Use of Laboratory Animals, and approved by Columbia University’s Institutional Animal Care and Use Committee. Monkeys were surgically implanted with a head post and recording cylinder using aseptic technique. Cortical area MT (left hemisphere in monkey D, right in monkey N) was initially targeted using structural MRI scans and confirmed using established physiological criteria prior to injection of virus.

Viral injections were performed while the animals were awake and seated in a primate chair. A glass microinjection pipette (115 μm outer diameter; Thomas Recording GmbH, Giessen, Germany) was affixed to a tungsten microelectrode (75 μm shank diameter; FHC, Inc., Bowdoin, ME) with cyanoacrylate to create a custom ‘injectrode’ which was passed through a transdural guide tube and advanced into area MT using a hydraulic microdrive. The metal connector at the back of the pipette was connected with flexible tubing to a Hamilton syringe loaded with 5-10 μl of virus (AAV8-CamKII-Jaws-KGC-GFP-ER2; titer = 5.9×10^12^ genomes/ml; UNC Vector Core, Chuong et al. 2014).

After locating a stretch of gray matter consistent with the known functional properties of MT, we advanced the injectrode to the deepest point of this stretch and began a series of injections. Using a syringe pump (Harvard Apparatus, Holliston, MA), we injected 0.75-1.0 μl at a rate of 0.05 μl/min at each of several locations spaced 400-500 μm apart. Each injection was followed by a 10 minute pause before slowly (5 μm/s) retracting the injectrode to the next site. This process continued until reaching the shallowest portion of the target region, resulting in 5-8 sites and a total of 4-7 μl injected along a given track. On subsequent days, the procedure was repeated in 1 (monkey N) or 3 (monkey neighboring grid holes (1 mm apart) for a total injected volume of 13 and 19 μl, respectively. We then waited at least 8 weeks before beginning experiments. The tissue remained viable and responsive to light for at least 12 months post-injection in monkey D and at least 9 months in monkey N. Some virus likely reached portions of nearby area MST, but all sites tested in the behavioral task fit the criteria for MT > MST, including RF size versus eccentricity, preference for slower dot speeds (<15-20°/s), and position relative to the white matter ventral to the sulcus.

To examine viral vector-mediated opsin expression histologically, we injected a third animal (monkey with the same procedure and identical vector stock used in monkeys N and D. Monkeys N and D remain actively contributing to other projects, thus precluding histological analysis of these animals. Monkey E received a total of 6.5 μl of vector delivered at 7 injection sites spaced 400 μm apart along a single penetration. These injections were made in the lateral bank of the intraparietal sulcus, rather than in MT, because of the constraint imposed by a previously existing recording chamber.

One month following injections, monkey E was euthanized with an overdose of pentobarbital and perfused transcardially with 4% paraformaldehyde followed by a gradient of sucrose in phosphate buffer (10, 20 and 30%). The brain was removed and cryoprotected in 30% sucrose. Coronal sections (50 μm) were cut on a sliding microtome and mounted onto slides. Transduced cells were first localized by inspecting native fluorescence signals in a series of sections spanning the intraparietal sulcus. Additional sections near the region of strongest expression were processed immunohistochemically using primary antibodies against GFP (Abcam 13970 RRID: AB_300798, 1:1000; Abcam, Cambridge, MA) and against the pan-neuronal marker NeuN (Millipore MAB377 RRID AB_2298772; MilliporeSigma, Burlington, MA) or the inhibitory neuron marker parvalbumin (Swant PV235 RRID AB_10000343 1:5000; Swant, Marly, Switzerland), and using secondary antibodies (Invitrogen Molecular Probes, Thermo Fisher Scientific, Waltham, MA): Alexa 594 (A21203 RRID: AB_141633, 1:400), Alexa 568 (A10042 RRID: AB_2534017, 1:400), Alexa 488 (custom, 1:400) and the nuclear stain DAPI (Invitrogen Molecular Probes D-21490, 1:5000) for visualization by epifluorescence microscopy. An upper bound on transduction selectivity for excitatory neurons was coarsely estimated by counting the number of GFP-positive, PV-positive and double-labeled somata in a single section imaged at 20X.

### Optrode design, site selection and photosuppression protocol

Photosuppression was achieved using a custom optrode of similar design to the injectrode described above: a tungsten microelectrode (75 or 100 μm shank diameter, ∼1 MOhm; FHC) glued to a optical fiber (Thorlabs UM22-100, core+cladding diameter = 110μm; Thorlabs, Newton, NJ). The fiber tip was sharpened as follows to reduce tissue damage and generate a broader light cone (Dai et al., 2014; Hanks et al., 2015). After stripping the fiber jacket, the tip was immersed in 48% hydrofluoric acid to a depth of 0.5-1.0 mm, with the polyimide coating intact to enhance smoothness and reproducibility of the resulting tip (’tube etching’; (Lambelet et al., 1998; Stöckle et al., 1999). The immersion took place within the narrow end of a standard 100 μl pipette tip containing the acid and sealed with Parafilm. After 30-45 minutes, the acid was rinsed off and the polyimide coating mechanically stripped, leaving 6-10 mm of bare cladding and a conical tip with full angle of approximately 15°. The sharpened fiber was then passed through the guide tube along with an electrode and the two were glued together using a tiny amount of cyanoacrylate, with the electrode tip positioned 300-500 μm ahead of the fiber tip. The source end of the fiber patch cable was connected to a 633 nm diode laser (LuxX+ 633-100, Omicron-Laserage Laserprodukte GmbH, Rodgau, Germany) under analog and digital control.

Prior to each session, we used a handheld power meter (Thorlabs PM160) to calibrate the total light power as a function of analog input to the laser controller. Given the known values for the beam half-angle in air (20-30°) and separation between fiber and electrode tips (300-500 μm), we estimated the irradiance of neurons near the electrode tip using the brain tissue light transmission calculator provided by the Deisseroth Lab, Stanford University (https://web.stanford.edu/group/dlab/cgi-bin/graph/chart.php). The goal (see below) was to use the minimum irradiance required to significantly reduce the firing rates of neurons within a small region of interest (300-500 μm diameter), taking advantage of the fact that irradiance in tissue falls off exponentially with distance. Thus, total light power was nearly always kept below 2 mW (16 mW/mm^2^ at a distance of 300 μm) and was typically between 0.3 and 1.0 mW (1.2-4.0 mW/mm^2^) during the discrimination task.

After advancing the optrode into area MT and pausing 20-40 minutes to allow the tissue to stabilize, we isolated single-unit (SU) or multi-unit (MU) activity using Plexon SortClient software (Plexon, Inc., Dallas, TX). RF size, position, and selectivity for motion direction and speed were assessed using briefly flashed, 99% coherence RDM stimuli while the animal fixated a central target. Candidate sites were mapped extensively before each behavioral session to ensure consistency of tuning/RF parameters across at least 200 and preferably 300+ μm of cortex. Responsiveness to red light was concurrently assessed by randomly interleaving laser trials (laser+RDM) with no-laser trials (RDM only), and sites were bypassed unless we observed a significant decrease in MU firing rate on laser versus no-laser trials (*t*-test, p<0.01) throughout the target area. A site was considered provisionally acceptable if (a) RF and tuning parameters remained relatively stable (Δ preferred direction < 45°) for at least 200 μm, (b) direction selectivity was sufficiently strong, with at least 2 standard deviations separating preferred and null (antipreferred) direction MU responses, and (c) activity on laser trials was reduced by at least 10%. Once we encountered an acceptable site, we attempted to position the optrode at the optimal depth with respect to the above considerations, and commenced the discrimination task. Twenty-seven sites were provisionally accepted (17 in monkey D, 10 in monkey N), but 4 sites in monkey D were rejected post-hoc after failing to show robust suppression of MU activity throughout the session.

On a random half of discrimination trials, including T_s_-present and T_s_-absent trials, red light was delivered throughout the visual stimulus epoch. The laser power profile was constant-on (square pulse), but in a majority of sessions was terminated with a 140 ms linear ramp-down to reduce the post-suppression burst (Fig. 3D and inset; Chuong et al., 2014). Laser onset began 20 ms after stimulus onset to partially account for visual response latency yet ensure that photosuppression began before the earliest feedforward visual inputs reached MT. Onset of the ramp-down began 20 ms after visual motion offset.

The fiber jacket and optrode shank were covered with opaque material such that no laser light was visible to the animal. Nevertheless, to rule out possible nonspecific effects of illumination, including tissue heating, we performed 8 control sessions in which the optrode tip was positioned either near the surface of the brain (N=2), or in the vicinity of the dorsal superior temporal sulcus but at least 2 mm outside the viral injection site (N=6). No behavioral effects of illumination were observed in these control sessions (Supplementary Fig. S6).

### Data analysis

We quantified the effect of photosuppression on behavioral choices using the logistic regression model:

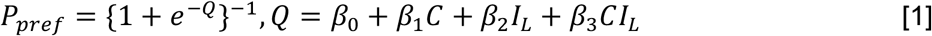

Where *P*_*pref*_ is the probability of a preferred-direction choice, *C* is signed motion coherence (positive = preferred direction, negative = antipreferred direction), and *I*_*L*_ is an indicator variable for the presence/absence of laser illumination. The biasing effect of photosuppression is expressed in units of coherence by the ratio *β*_5_/*β*_0_, and its effect on discrimination sensitivity (slope) is captured by *β*_8_. The coefficients were fit by maximum likelihood estimation, with standard errors (SEs) estimated as the square roots of the diagonal elements of the inverse of the Hessian matrix. SEs were used to calculate *t*-statistics and corresponding p-values to test the null hypothesis that a given *β*_9_ = 0. To quantify the difference in sensitivity when T_s_ was available versus unavailable (Fig. 1C), we replaced *I*_*L*_ in Eq. 1 with an indicator variable for the presence/absence of T_s_. For display purposes, smooth curves in Figs. 4C, 5A+B, and 7A+B were generated using the simpler model:

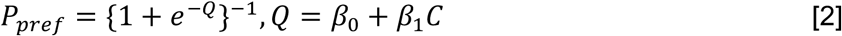

fitted separately to laser and no-laser trials.

The effect of photosuppresion on confidence was quantified by fitting the probability of a sure-bet choice (*P*_*sb*_) with the Gaussian function:

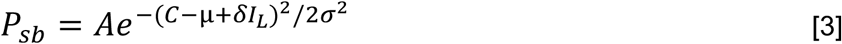

Where *C* is signed coherence, *I*_*L*_ is the indicator variable for the laser, and *A, μ, σ*, and δ are free parameters. The parameter δ captures the photosuppression-induced lateral shift of the Gaussian in units of coherence.

For each laser trial, we estimated the degree of Jaws-mediated suppression of neural activity as follows. Multi-unit spike events were counted from 60 ms after RDM onset until 60 ms after RDM offset, then converted to spike rate (*R*_*laser*_) by dividing by the stimulus duration. The fractional change in spike rate (*ΔR*) was then computed by comparing each *R*_*laser*_ with the mean spike rate of a matching set of no-laser trials 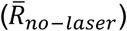, defined as being of the same session, motion direction, coherence, and quartile of stimulus duration:

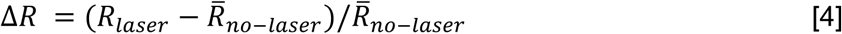

Behavioral effects as a function of Δ*R* (e.g., Fig. 4B) were computed by selecting laser trials with a given range of Δ*R* and comparing them to all no-laser trials across all sessions, using Eqs. 1 and 3 for choice and confidence, respectively. The traces in Fig. 4B were mapped out by repeating this procedure for each of many sliding windows of trials sorted by *ΔR*, where the window width was 2100 trials (approximately 1/5 of laser trials, i.e. a sliding quintile) and the step size was 10 trials. Similarly, Figures 6A and 6B were constructed by sorting trials by the variable on the abscissa (conditioned on two other post-hoc variables; see below) and computing behavioral effects using a sliding quartile window.

Having observed variation in the size of behavioral effects as a function of three main explanatory variables—fractional change in spike rate on laser trials (Δ*R*), trial number within session (*T*), and stimulus/laser duration (*D*)—we performed post-hoc statistical tests of these relationships using binomial generalized linear models (i.e., logistic regression). Each model consisted of three predictors, plus all 2-way interaction terms. The predictors were: signed motion coherence (*C*), the indicator variable for presence of the laser (*I*_*L*_), and an indicator variable defining a median split for one of the three variables mentioned above (Δ*R, D*, or *T*). For example, the model to test the influence of stimulus duration *D* on photosuppression-induced choice effects was specified by:

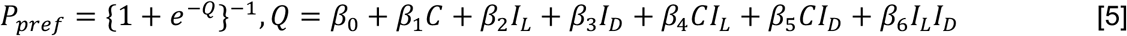

Where *I*_*D*_ = 1 if D < 300 ms (the median duration) and 0 otherwise. The model for Δ*R* substituted *I*_*ΔR*_ (1 if Δ*R* < −0.35) in place of *I*_*D*_, and for trial number the corresponding variable was *I*_*T*_ (1 if T < 519). Since the relationship between trial number and the confidence effect (Fig. 6A, blue) was found to be non-monotonic, we used a smaller cutoff value of T < 300 to define *I*_*T*_ for that test. For each model, the strength of the interaction between the given explanatory variable and the biasing effect of photosuppresion is captured by *β*_6_and its associated p-value.

A similar strategy was used for the effects on confidence (replacing *P*_*pref*_ with *P*_*sb*_ in Equation 5 and its counterparts for Δ*R* and *T*), except that the models were fit separately for trials with *C* < 0 and *C* > 0. This piecewise-logistic approach achieved adequate fits because each side of the bell-shaped relationship between *P*_*sb*_ and *C* is well approximated by a sigmoid, albeit of opposite slope. This yielded two *β*_6_ coefficients and corresponding p-values, and the test was deemed significant if either were less than 0.05 after Bonferroni correction (multiplication by 2). To better isolate the effects of trial number, all GLM analyses, as well as the sliding-window plot in Fig. 6A, included only trials with ΔR < −0.25 and D < 300 ms. Similarly, the GLMs for duration (and Fig. 6B) were limited to only trials with ΔR < −0.25 and T < 500.

To compare neuronal direction selectivity across different subsets of trials—e.g., short vs. long trials (Fig. 7)—we used a standard approach to calculate neural thresholds (Britten et al., 1992). For each trial in a given session, the MU spike rate was normalized to the mean spike rate for 51.2% coherence motion in the preferred direction during that session. Normalized spike rates were then pooled across sessions, grouped according to desired criteria (duration, trial number within session, etc.), and the distributions of spike rates for preferred-vs. antipreferred-direction motion were compared for a given coherence level using ROC analysis. This quantified the ability of an ideal observer to discriminate motion direction based on the neural responses. The performance of the ideal observer (percent correct as a function of coherence) was fit with a cumulative Weibull distribution (Quick, 1974) and the *α* parameter (coherence level generating 82% correct) was taken as the neuronal threshold.

---

**Supplementary Figure S1.**
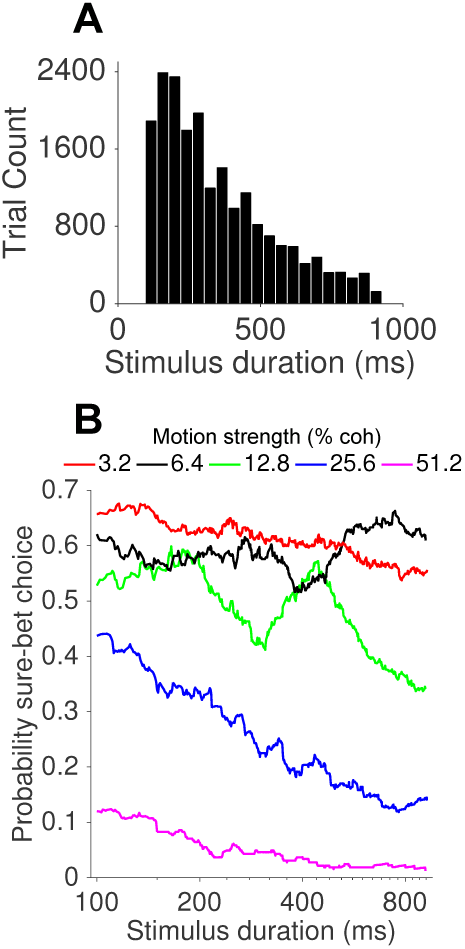
**(A)** Histogram of the duration of visual stimuli (and photosuppression, on laser trials, not including the ramp-down). Duration on each trial was drawn from a truncated exponential distribution with range 95-925 ms and median 300 ms. **(B)** The probability of sure-bet choices as a function of stimulus duration and motion strength.

**Supplementary Figure S2.**
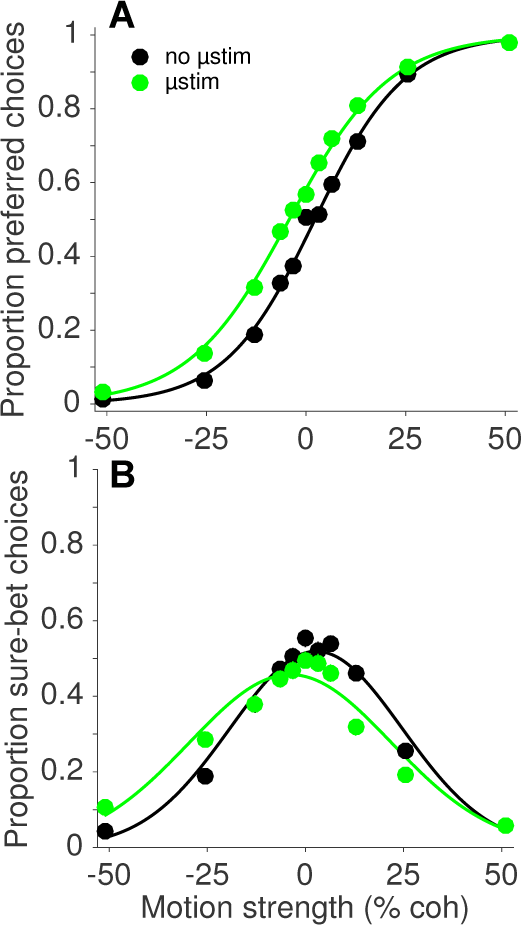
Electrical microstimulation (μStim) data from one monkey used in a previous study (Fetsch et al., 2014) who also participated in the present study, showing **(A)** a leftward shift of the choice function on μStim trials (green) and **(B)** a corresponding shift in the confidence function.

**Supplementary Figure S3.**
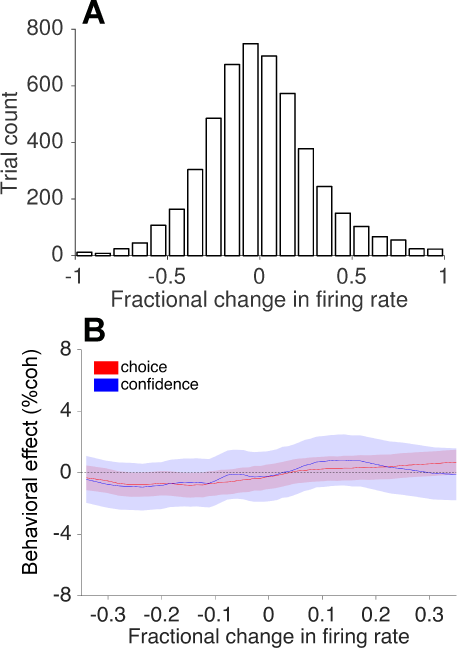
Control analysis (mimicking Fig. 4A+B) testing whether variability in responsiveness in the absence of photosuppression can explain shifts in choice and confidence functions. **(A)** Distribution of fractional ‘change’ in firing rate for each no-laser trial relative to the mean of trials from the corresponding session and trial type. **(B)** Sliding-window analysis of randomly assigned ‘sham-laser’ trials sorted by the abscissa value in A, showing the size and direction of shifts in choice and confidence as a function of variability in firing rate.

**Supplementary Figure S4.**
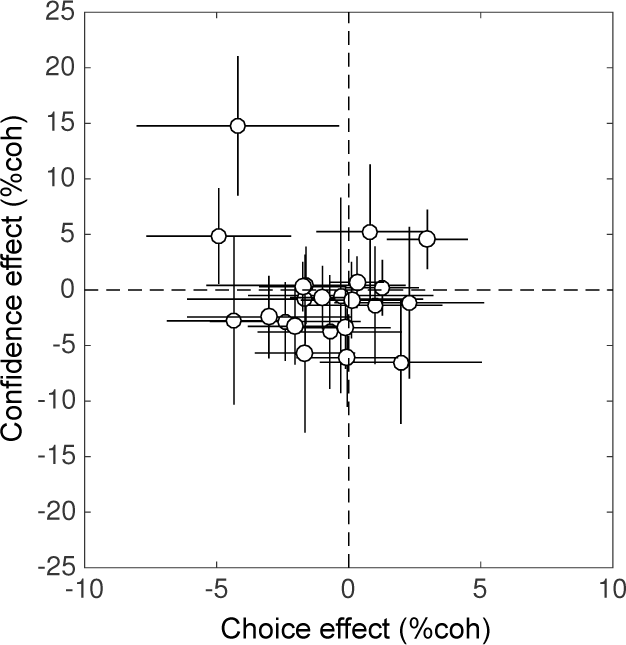
Effects of photosuppression on choice (abscissa) and confidence (ordinate) based on all trials in each session (N=23, mean = 872 trials per session, S.D. = 487 trials). Negative values indicate shifts in the predicted direction based on the preferred direction of the neurons. Error bars are ± SEM.

**Supplementary Figure S5.**
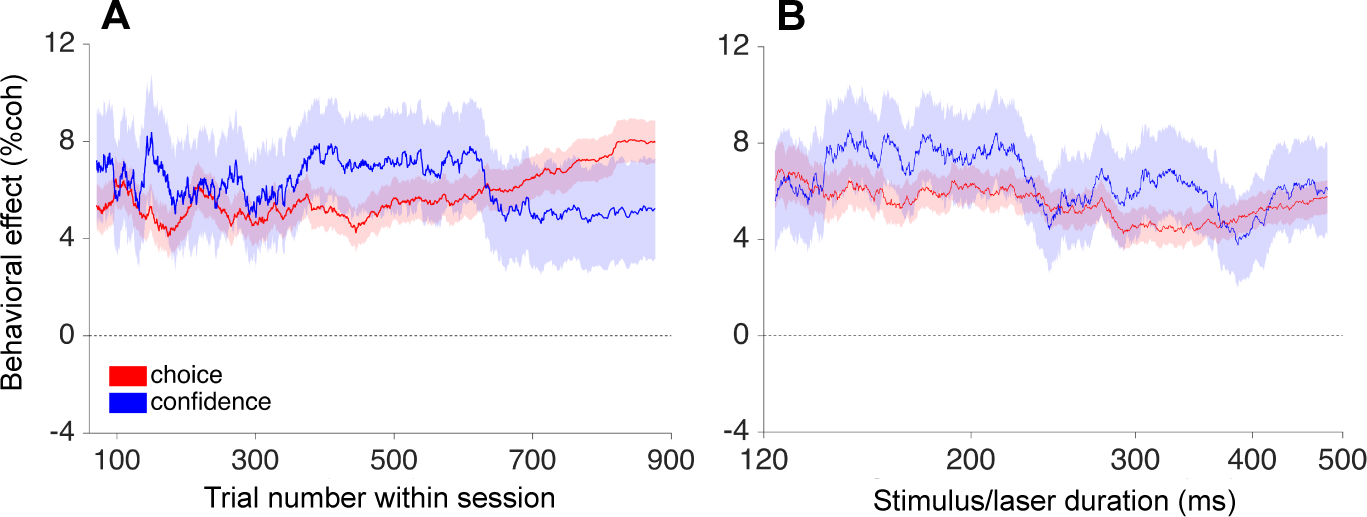
**(A,B)** Results of the same analyses shown in Fig. 6, applied to a previously obtained μStim dataset. The compensatory effects seen with photosuppression were absent for μStim, which produced relatively consistent effects on choice and confidence over the course of the session (A) and on trials of different duration (B).

**Supplementary Figure S6.**
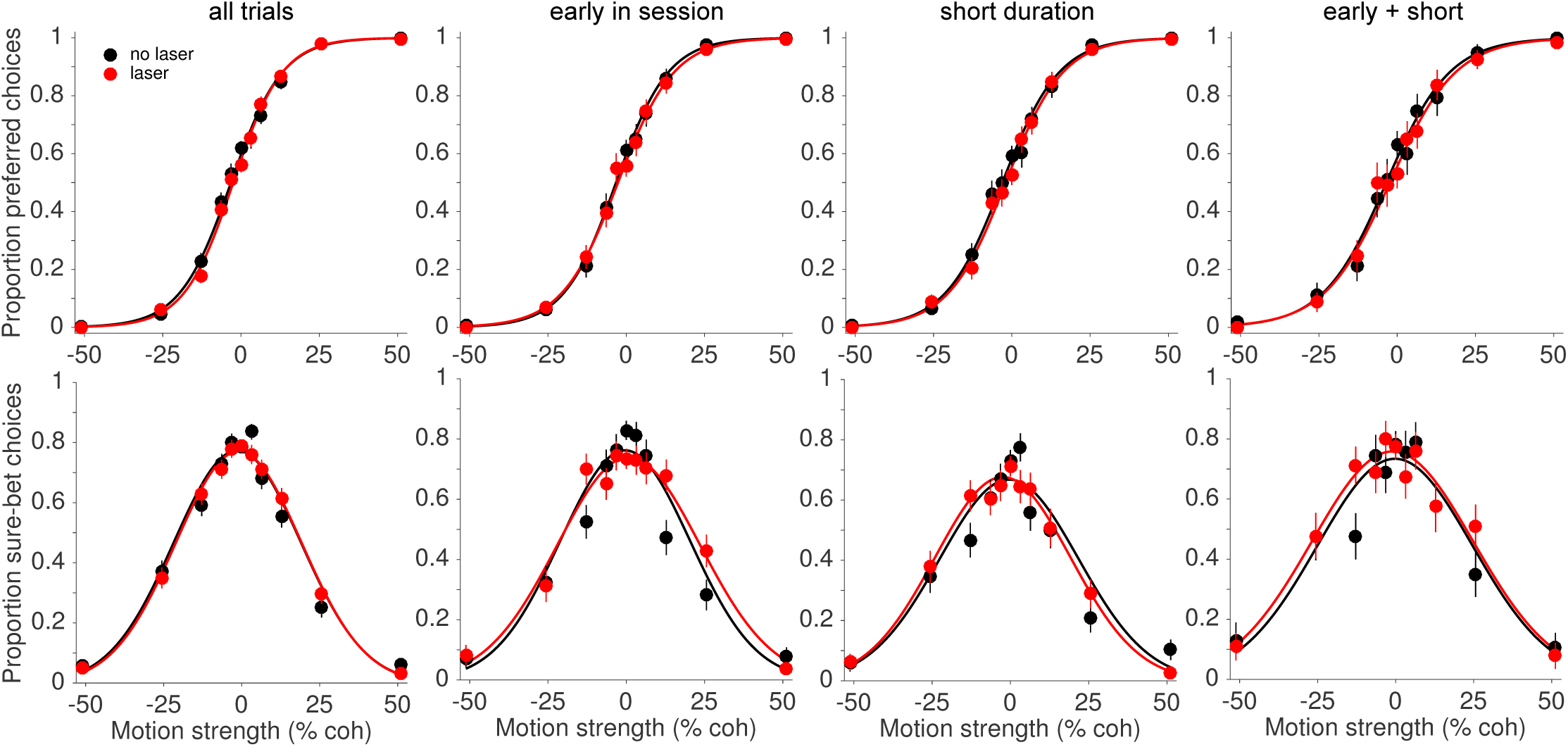
Control experiments to rule out nonspecific (non-opsin-mediated) effects of laser illumination. The optrode was positioned either just inside the guide tube or deep within the brain but away from the virus injection site. Left panel, N=7491 trials across 8 sessions. Subsequent panels include only trials early in the session, of short stimulus duration, or those meeting both conditions.

